# Increased TSH-producing cells in the pituitary gland of Pax6 haploinsufficient mice

**DOI:** 10.1101/2020.04.06.028282

**Authors:** Kenji K. Johnson, James D. Lauderdale

## Abstract

Aniridia is a congenital condition characterized by absence of iris and is caused by a semidominant mutation in the transcription factor encoded by the *PAX6* gene. Although ocular phenotypes of this disorder are well characterized, recent studies report that individuals with aniridia have a higher propensity for obesity, infertility, polycystic ovarian disease, and severe eczema compared to their *Pax6*-normal siblings. These symptoms collectively suggest an underlying endocrine disturbance related to haploinsufficient levels of *Pax6.* In mice, during development, *Pax6* expression in the pituitary gland begins at E9.0 in the primordial anterior pituitary gland (Rathke’s Pouch). This expression becomes restricted to the dorsal anterior pituitary by E11.5, but is expressed throughout the anterior lobe by E14.5, and remains through adulthood. It is possible that a reduction in *Pax6* could result in a change in pituitary hormone levels or cell numbers, which may explain symptoms experienced by aniridics. Using the *Small eye* mouse model, we find that *Pax6* reduction results in a decrease in GH-producing cells and an increase in TSH-producing cells in neonate mice, with the TSH increase continuing into adulthood. Adult *Pax6* haploinsufficient mice also have an increase in anterior pituitary volume and weigh significantly less than their wild-type littermates. Furthermore, we show that the increase in TSH-producing cells leads to an increase in thyroxin (T_4_) in mutant mice, although tri-iodothyronine (T_3_) levels remain unchanged. These findings present a new role for *Pax6* in the endocrine system, which serves to refine our current understanding of *Pax6* in endocrine development and maintenance and provides new avenues for investigating endocrine-related symptomatology in aniridia.

## Introduction

*Pax6* is a highly conserved transcription factor that is expressed in the developing central nervous system (CNS) where it is required for proper development of tissues such as the eye, forebrain and hindbrain, and spinal cord (1–3). In addition, *Pax6* is also expressed in the pancreas, specifically the islets of Langerhans, the L and K cells of the small and large intestine, and the pituitary gland (3–7). Homozygous mutations in *Pax6* results in lack of eyes and deformed nasal cavity while a heterozygous loss of function mutation in humans causes a condition known as aniridia (8–10).

Aniridia is a rare, congenital disorder that occurs in approximately 1 in 50,000 to 100,000 live births and is mainly characterized by complete or partial absence of the iris (11,12). In addition to the iris, it is well established that it also results in vascularization of the corneas, glaucoma (leading to vision loss), and cataracts (13–15). Although the eye defects associated with *Pax6* and aniridia have been studied, less is known about the systemic effects of a deficiency of *Pax6.* Recently, people with aniridia have self-reported a higher propensity for systemic symptoms including obesity, polycystic ovarian disease, infertility, and severe eczema (16). Based on previous studies (20), we hypothesized that these symptoms may be due to a perturbation of the endocrine system, specifically a change in cell numbers within the anterior pituitary gland.

The pituitary gland consists of three distinct parts: the posterior and anterior lobes and, in rodents, an intermediate lobe. While the posterior lobe develops from an evagination of the infundibulum of the hypothalamus, the intermediate and anterior lobes are derived from an invagination of the oral ectoderm into its primordial structure, Rathke’s Pouch (17,18). The oral ectoderm begins to thicken and invaginate around E7.0 in mouse development. This invagination continues until the full development of the Rathke’s Pouch by E10.5 and is fully separated from the oral roof ectoderm by E12.5 (19). Upon this separation, the five cell types of the anterior pituitary gland characterized by hormone production begin their differentiation through E18.5 in a dorsal to ventral pattern: adrenocorticotropin hormone (ACTH), growth hormone (GH), prolactin (PRL), thyroid stimulating hormone (TSH), follicle stimulating hormone (FSH), and luteinizing hormone (LH).

Coincidentally, in addition to its expression in the anterior neural ridge and oral ectoderm beginning at E8.0, *Pax6* is also expressed dorsally in the Rathke’s Pouch between E10.5 and E12.5 and throughout the anterior pituitary gland at least through E18.5 in the developing mouse (3). Furthermore, previous studies have shown that in the complete absence of *Pax6,* there is a decrease in the more dorsal cell types GH and PRL and an increase in the ventral cell types TSH and LH (20,21). Although these studies specifically addressed homozygous loss of function of *Pax6*, if these changes in cell numbers are also present in the heterozygous condition, it might explain some of the systemic symptoms experienced by people with aniridia. Further, recent studies have shown an association between *PAX6* mutation in humans and a disruption in pituitary hormone activity (22–24).

Here, we used the mouse model for aniridia known as *Small eye* to investigate a potential change in hormone cell numbers of the anterior pituitary gland in mice heterozygous null for *Pax6.* We show that *Pax6* reduction results in a decrease in GH-producing cells and an increase in TSH-producing cells in neonate mice, with the TSH increase continuing into adulthood. This increase in TSH-producing cells leads to an increase in thyroxin (T_4_) in adult mutant mice, although tri-iodothyronine (T_3_) levels remain unchanged. These data support the hypothesis that *Pax6* plays a role in the adult endocrine system.

## Materials and Methods

### Animals

The mice used for this study were maintained as a *Pax6^Sey-Neu/+^* colony on a majority CD1 genetic background. Wild-type (*Pax6*^+/+^) littermates were used as controls. The genotype of each animal was determined by PCR as previously described (25). Both males and females were used. All experiments involving mice were conducted in strict accordance with the National Institutes of Health Guide for the Care and Use of Laboratory Animals and were performed with approval and oversight of the University of Georgia Institutional Animal Care and Use Committee.

### Histology and immunohistochemistry

To rapidly and uniformly preserve brain tissue after euthanization, adult mice were perfused with 1x phosphate-buffered saline (PBS) and 4% paraformaldehyde/PBS (PFA). Tissues obtained from neonatal (P0) animals and dissected adult pituitary glands were preserved by immersion in 4% PFA at 4°C overnight, rinsed in 1x PBS, and then dehydrated stepwise through a graded series of 50, 70, 90, 96 and 100% ethanol, equilibrated with xylene, and embedded in paraffin using a Tissue Tek apparatus (Miles, Ekhart, USA). Serial sections were cut using a rotary microtome at 8 μm, mounted onto slides (Superfrost/Plus; Fisher Scientific, Pittsburg, PA), and dried at 37°C overnight. Tissue sections were deparaffinized by two rinses in xylene followed by rehydration in decreasing concentrations of ethanol, with a final rinse in tap water. Sections used for histology were stained with Mayer’s hematoxylin (Sigma) and eosin solution.

Sections used for indirect immunofluorescence were bleached by incubation in aqueous 3% hydrogen peroxide for 10-20 minutes, and subjected to citrate buffer antigen retrieval. Briefly tissue sections were placed into a boiling sodium citrate solution (10mM Citric Acid, 0.05% Tween 20, pH 6.0) for 30 minutes, removed, and allowed to cool for 30 minutes at room temperature. Sections were then rinsed in PBS and blocked for 30 minutes in a solution of 1% bovine serum albumin (BSA; Fisher Scientific, catalog BP1600-100), 5% normal donkey serum (Sigma-Aldrich, catalog D9663) in PBS. Overnight primary antibody incubation was performed in blocking solution at 4°C in a humidified chamber. Unless otherwise stated, primary antibodies were obtained from the National Hormone and Peptide Program (NHPP, Harbor-UCLA Medical Center, Torrance, CA). Primary antibody identities and dilutions were as follows: rabbit anti-ACTH (AFP156102789Rb, 1:100), rabbit anti-GH (AFP5641801Rb, 1:1000), rabbit anti-TSH (AFP1274789Rb, 1:100), rabbit anti-LH (AFP571292393Rb, 1:100), guinea pig anti-FSH, and guinea pig anti-PRL. After removal of the primary antibody, tissues were rinsed three times with PBS for 5 minutes each, blocked for 10 minutes, and incubated for 30 minutes at room temperature with a 1:1000 dilution of donkey anti-rabbit IgG (H+L) secondary antibody conjugated to Alexa Fluor^®^ 647 (ThermoFisher-Invitrogen, catalog A-31573) in blocking solution. After removal of the secondary antibody, the tissue sections were rinsed several times with PBS, the nuclei were labeled with a 1:10,000 dilution of DAPI (4’,6-Diamidino-2-phenylindole; Sigma, D9542) in PBS, and then coverslipped with EMS-Fluorogel.

### Imaging of immunolabeled sections

Specific signals were visualized using either standard fluorescence microscopy using a Zeiss Axio Imager.D2 or laser scanning confocal microscopy using a Zeiss LSM 510 Meta Confocal Microscope and images acquired using Zeiss image acquisition software. Between 3 and 5 pituitaries were analyzed for each stage and genotype; representative images are shown. N-values for each experiment are provided in the text and figure legends.

### Cell counts

For cell counts made of the pituitary glands from neonates, the pituitary glands from *Pax6^+/+^*, *Pax6^+/−^* and *Pax6^−/−^* mice at P0 were serially sectioned in their entirety and immunolabeled for ACTH, GH, TSH and LH and counter stained with DAPI as described above. For each of these hormone-producing cell types, the total numbers of specifically immunolabeled cells were counted in every other section and the numbers for each pituitary totaled. The average number and standard deviation (SD) of immunolabeled cells per genotype were determined using three or four animals per group. Statistical comparison between genotypes was performed by analysis of variance (ANOVA).

### RNA *in situ* hybridization

RNA *in situ* hybridization was performed as previously described (26), with the exception that the proteinase K digestion was omitted, on tissue sections prepared from 7 months old adult wild-type mice. Sense and antisense digoxigenin-labeled RNA probes were prepared from a BglII-digest of the pMPX2-1 mouse *Pax6* cDNA clone (25) using a DIG RNA labeling kit (Roche). Hybridization and stringent posthybridization wash steps were performed at 65°C.

### RT-PCR

Total RNA from whole eyes, pituitary gland, lung and heart from a 9 months old wildtype adult was prepared using TRIzol reagent (ThermoFisher-Invitrogen, catalog 15596026) following the manufacturer’s recommended conditions. Total RNA was then treated with the Turbo DNA-freeTM Kit (ThermoFisher-Ambion, catalog AM1907) following the standard protocol to remove potential DNA contamination. The treated RNA was then reverse transcribed (SuperScript Double-Stranded cDNA Synthesis Kit), and the resulting cDNA was amplified by PCR using a mouse *Pax6* primer pair designed by PrimerBank (27–30). This primer pair (PrimerBank ID: 7305369a1) generates a 285 bp amplicon spanning exons 5a - 7. PCR conditions as previously described (31).

### Western Blot

Western analysis was performed on protein lysates prepared from whole eyes, pituitary glands, lung and heart dissected from adults (7 months old). Lysates were prepared by homogenization of tissues in RIPA buffer (10mM Tris-HCl, 150mM NaCl, 1mM EDTA, 1% NP-40, 0.1% SDS, 10% glycerol, 1mM PMSF, 1mM EGTA) in ice. Homogenized samples were centrifuged at 13,000 rpm for 5 minutes at 4°C, and the supernatant collected. Bradford Protein assay method was used to determine the protein concentration using bovine serum albumin (BSA) as the standard (Bio-Rad catalog 500-0006). 1-20μg of tissue protein to be analyzed were combined with equal parts 2X Laemmli sample buffer (Bio-Rad, catalog 161-0737) containing 10% β-mercaptoethanol, and then boiled for 15 minutes, loaded onto a reducing-denaturing SDS-polyacrylamide gel (10-12% resolving, 4% stacking), and one-dimensional electrophoresis was carried out for 4 hours at 75-100 volts in a Tris-glycine-SDS (Bio-Rad, catalog 161-0732) running buffer. Proteins were transferred to a 0.45μm nitrocellulose membrane (Bio-Rad, catalog 162-0115) in Tris-glycine transfer buffer containing 20% methanol cooled to 4°C. After transfer, the membrane was blocked overnight at 4°C using 5% non-fat dry milk (Bio-Rad, catalog 170-6404) in Tris-buffered saline pH 7.3, 0.1% Tween 20 (TBSTw) with agitation. Blots were incubated for 1 hour at room temperature with a 1:2000 dilution of a rabbit anti-*Pax6* primary antibody directed against the C-terminus of the protein (Covance, catalog PRB-278P) in blocking solution with gentle agitation. Following primary antibody incubation, membranes were rinsed several times in 1xTBSTw and incubated with goat anti-rabbit HRP-conjugated secondary antibody (1:20,000; Bio-Rad, catalog 179-5046; or 1:10,000; Santa Cruz, catalog sc-2004) in 5% non-fat dried milk for 1 hour at room temperature with gentle agitation. After removal of the secondary antibody, the membranes were rinsed several times in TBSTw. For signal detection, Immuno-Star Western C Kit (BioRad catalog 170-5070) was used, followed by manual autoradiograph development. To determine protein loading, membranes were subsequently stripped (32), blocked in 10% milk in 1xTBSTw overnight at 4°C and incubated with rabbit anti-GAPDH primary antibody (1:1000, abcam, catalog ab9495) and processed as described above.

### Flow Cytometry

Freshly dissected pituitary glands from adult mice (6-8 months of age) were then extracted and mechanically dissociated in 1x PBS followed by fixation in 4% PFA. Fixed cells were then permeabilized in 90% methanol for 30 minutes on ice, followed by several washes in 1xPBS, leaving 100ul of solution in each wash. Cells were then stained according to established protocols for immunofluorescence staining of cells for flow cytometry (33,34) using the same primary antibodies listed above for immunofluorescence and goat-anti rabbit allophocyan (APC) for secondary antibody. APC fluorescence was detected through 665/20 filter with logarithmic amplification. Cells were initially gated on a scatter plot and a FSC pulse width vs. FSC peak plot to eliminate debris and doublets, respectively. 30,000 events were collected for each sample. Acquisition was performed using a CyAn ADP Analyzer (Beckman Coulter, Hialeah, Florida) and data analyzed using FlowJo software version 9.3.1 (Treestar, Inc., Ashland, Oregon).

### Weight data

To obtain weight data, *Pax6^+/−^* mice and their wild type littermates were euthanized by carbon dioxide followed by cervical dislocation and weighed immediately after euthanasia. The mice were 6 to 9 months of age. For each genotype 76 male mice and 79 female mice were weighed. For both males and females, equal numbers of wild-type and *Pax6^+/−^* mice were weighed at each age (males/females, 6 months: 35/33; 7 months: 14/8; 8 months: 13/6; 9 months: 13/32). Data was analyzed by Student’s t-test.

### Radioimmunoassays

Blood serum was isolated from individual adult *Pax6^+/−^* mice and wild-type littermates (6 males and 4 females per genotype, 4 months old) using BD serum isolation tubes. Samples were then frozen and sent to Michigan State University endocrine diagnostic labs to measure concentration of total T_3_ and T_4_. Data was analyzed by Student’s t-test.

## Results

### *Pax6* is expressed in the adult anterior pituitary gland

Although previous studies have reported on *Pax6* expression in the developing pituitary gland (3,20), there have been no reports of *Pax6* expression in the adult pituitary gland. Therefore, we examined adult pituitary glands from wild-type mice to determine if *Pax6* expression continues throughout adulthood. The mouse *Pax6* gene generates several different mRNAs and encodes for three protein isoforms (25). Using a probe designed to detect all *Pax6* transcripts, we found by RNA *in situ* hybridization that *Pax6* is expressed in the adult anterior pituitary but not in the intermediate or posterior lobes (Figure 1A) as was previously reported in the E18.5 embryo (3). Comparison of the RNA *in situ* hybridization signal observed in pituitary to eye tissue suggested that there was a much lower level of *Pax6* expression in the pituitary compared to cells in the retina (data not shown). To test this, reverse transcription polymerase chain reactions (RT-PCR) were performed on total RNA prepared from whole eye and pituitary under semi-quantitative conditions (Figure 1B, data not shown). *Pax6* transcripts were more abundant in total RNA prepared from whole eye than from pituitary. Although the RT-PCR experiments only capture bulk expression and will be strongly influenced by the numbers of cells that express the gene compared to those that do not within the different tissues, this data when combined with the RNA *in situ* data suggests that *Pax6* is expressed more or less uniformly by cells in the anterior pituitary and that the level of expression per cell is lower than those of the retina. Western analysis revealed the presence of the 46 kDa canonical PAX6 protein and the 48 kDa alternatively spliced exon 5a isoform (PAX6+5a) in the pituitary, albeit at lower levels than in the eye (Figure 1C). Together these results demonstrate that *Pax6* is expressed in the anterior pituitary gland in adult mice and suggest that, in addition to its role in the development of the pituitary gland, *Pax6* also likely plays a maintenance role in the adult anterior lobe.

**Figure 1:**
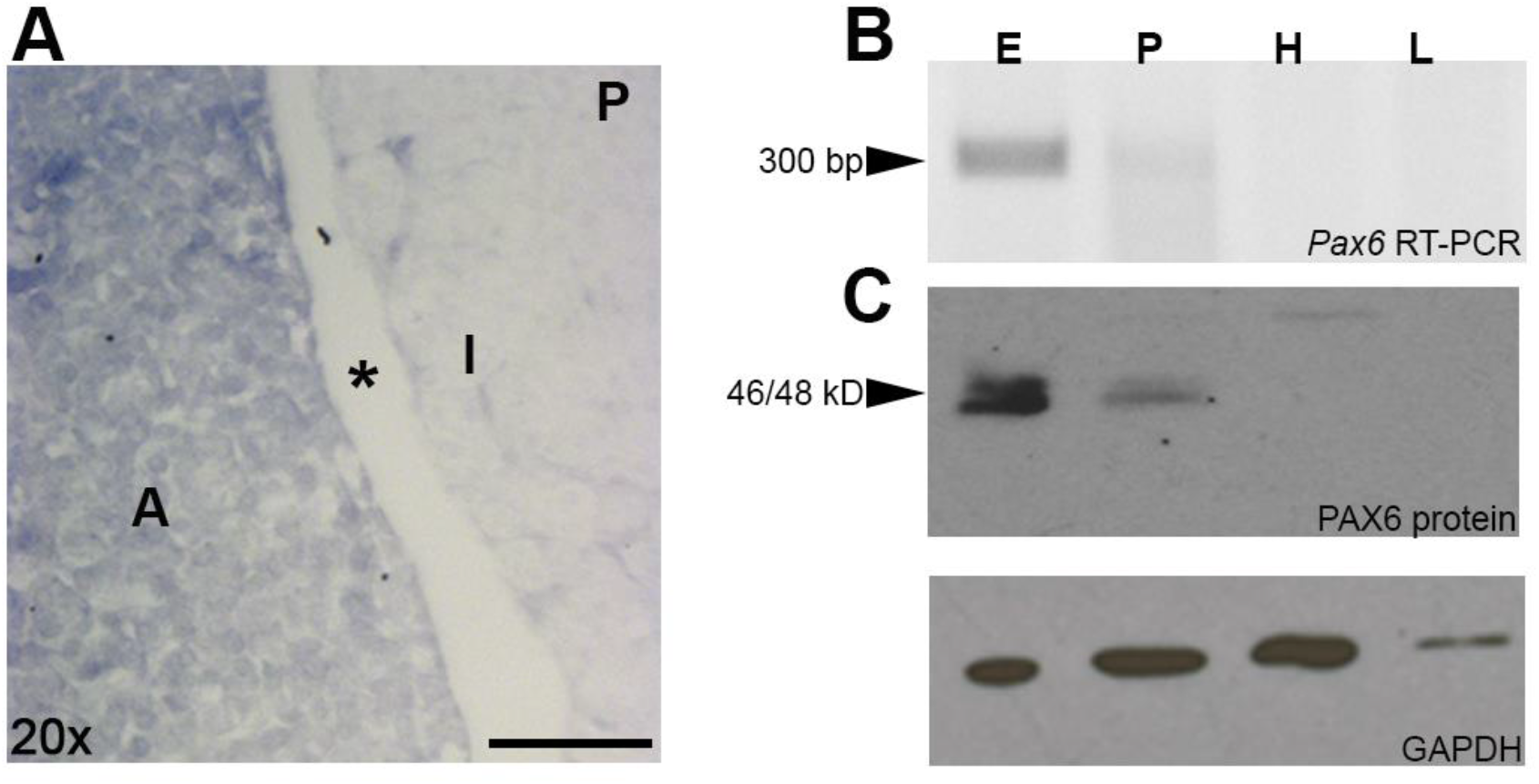
*Pax6* expression persists at low levels in adult pituitary glands. (A) Low levels of *Pax6* mRNA present in anterior pituitary gland as shown by RNA in situ hybridization. Abbreviations: Anterior pituitary (A), intermediate pituitary (I), posterior pituitary (P); scale bar = 100μm. (B) *Pax6* transcripts containing the alternatively spliced exon 5a were detected in total RNA prepared from the eyes (E) and pituitaries (P) of adult mice, but not heart (H) or lung (L). (C) Western blot shows presence of full-length PAX6 protein with (48 kDa) and without exon 5a (46 kDa) in lysates prepared from eye and pituitary, but not heart or lung. *In situ* hybridization and Western analyses were performed on tissues collected from 7 months old mice. RNA for the RT-PCR experiments was collected from 9 months old mice.

### Pax6 is required for normal morphological pituitary development

*Pax6* is known to be required for pituitary development. In *Pax6* homozygous null mice, there is a significant decrease in somatotropes and lactotropes in the anterior pituitary (20,21); however, potential modifier effects have been observed (21) and the net effect of *Pax6* dosage on pituitary development in neonates has not been reported. To provide a starting point for assessing potential changes in the adult pituitary of the *Pax6^+/−^* allele used the current study, we first examined by histology the pituitaries of neonate littermates at birth (Figure 2). Because mice homozygous null for *Pax6* die at birth, this was the oldest developmental time point at which we could compare pituitary morphology between *Pax6^+/+^, Pax6^+/−^* and *Pax6^−/−^* animals. Using the trigeminal ganglion as an anatomical landmark, we examined histological sections cut through the middle of the pituitary for each genotype. The H&E stained pituitary tissue sections from *Pax6^+/−^* neonates appear comparable to those from *Pax6^+/+^* neonates (compare Figure 2B to A). For both genotypes, the three lobes of the pituitary are clearly identifiable. The posterior lobe is located directly below the third ventricle of the brain and is adjacent to a clearly delineated intermediate lobe and the lumen separating it from the anterior lobe (Figure 2A,B). The lobes appear comparable in size (Figure 2A,B; data not shown). In contrast, sections cut through the *Pax6^−/−^* gland showed significant differences in the presumptive posterior pituitary lobe and the presumptive intermediate lobe was not distinct (compare Figure 2C to A). Notably in the null mutants, the cells in the presumptive posterior lobe of the pituitary exhibit histological characteristics similar to the anterior lobe, but do not express anterior lobe cell type markers (see Figure 4). The anterior lobe of the pituitary in *Pax6^−/−^* neonates was histologically more comparable to the anterior lobe present in the other genotypes, but appeared to be thicker in the area directly below the posterior lobe than was observed in either wild-type and *Pax6^+/−^* littermates (compare Figure 2C to A,B). These results demonstrate that the histological development of the pituitary is comparable between *Pax6^+/+^* and *Pax6^+/−^* mice at P0 and establish a baseline for the adult and cellular studies.

**Figure 2:**
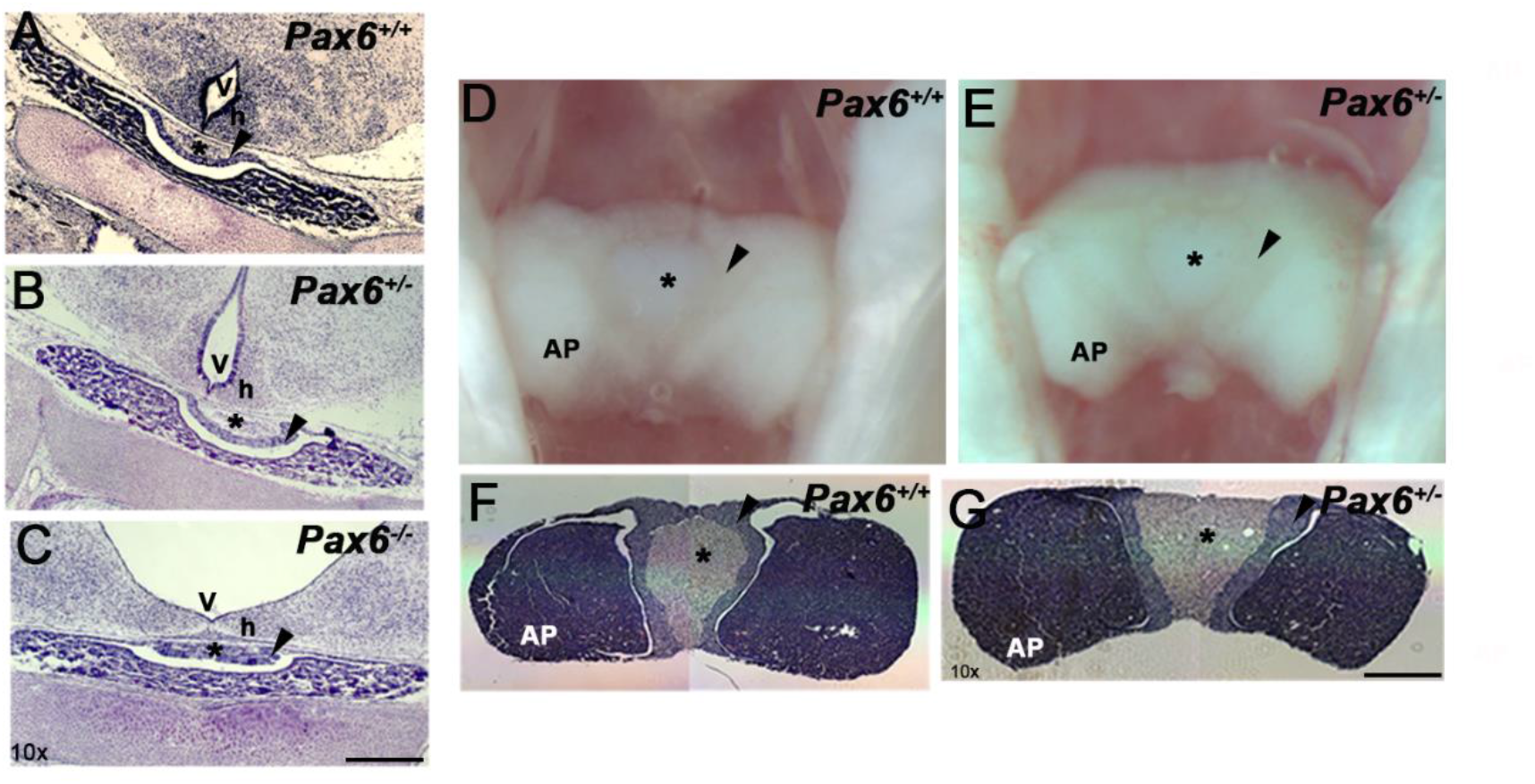
*Pax6* is required for normal pituitary development. Histological comparison of the pituitaries in *Pax6^+/+^* (A), *Pax6^+/−^* (B), and *Pax6^−/−^* (C) littermates at P0 age. The H&E stained pituitary tissue sections from *Pax6^+/−^* neonates appear comparable to those from *Pax6^+/+^* neonates, while those from *Pax6^−/−^* neonates show differences in the presumptive posterior (asterisk) and intermediate (arrowhead) pituitary. The enlarged ventricle in *Pax6^−/−^* neonates is due to aberrant brain development. AP, anterior pituitary; v, third ventricle; h, hypothalamus; asterisk denotes position of posterior pituitary, arrowhead denotes intermediate pituitary. Scale bar = 100μm. N=3 for each genotype. For D-G, comparison of pituitaries in *Pax6^+/+^* (D,F) and *Pax6^+/−^* (E,G) littermates at 8 months of age. (D,E) pituitaries *in situ.* (F,G) H&E stained transverse sections cut through the middle of the pituitary. The pituitary glands were comparable between genotypes. N=6 animals, 3 male and 3 female, for each genotype; animals were 6-8 months of age. AP, anterior pituitary; asterisk denotes posterior pituitary; arrowhead denotes intermediate pituitary. Scale bar = 100um.

### *Pax6^+/−^* pituitaries are morphologically normal but enlarged in adult mice

Since *Pax6* is semidominant and, in some organs such as the eye, the reduction in functional protein results in a change in morphology, we thought it was plausible that the structure and morphology of the mutant pituitary glands might also be affected. At both the gross and histological level, the pituitaries in adult *Pax6^+/−^* mice appeared comparable to those in wild-type littermates (Figure 2 D-G). The anterior, intermediate and posterior lobes could be clearly identified both *in situ* and in histological sections.

Casual inspection of the pituitaries *in situ,* in dissected whole tissues, and in sectioned tissues however suggested that the anterior lobe of the pituitaries from the *Pax6^+/−^* mice were slightly, but noticeably, larger than those from wild-type littermates. To avoid possible confounds associated with measuring freshly dissected tissues or changes in size associated with fixation and histology, we utilized MRI as a means of determining pituitary volume in live adult *Pax6^+/−^* mice and wild-type littermates. However, no difference in the size of the pituitary gland was observed by MRI (data not shown). Collectively, these data show that although there is no gross and significant difference in morphology of the pituitary glands of heterozygous mutant mice at birth, there may be a slight increase in the size of the anterior lobe during adulthood.

### Expression of endocrine cells in neonatal mice

In mice, pituitary hormone cell types develop in a dorsal to ventral pattern, with corticotropes (ACTH) dorsal-most, followed by somatotropes (GH) and lactotropes (PRL) in the intermediate position, thyrotropes (TSH) in the ventral-intermediate position and gonadotropes (FSH and LH) ventral-most. By E17.5, all cell types are present and have begun to take on their adult locations. Previous studies reported a role for *Pax6* in the dorsal to ventral patterning of the anterior pituitary gland, where the complete absence of *Pax6* resulted in a dorsal expansion of ventral cell types (gonadotropes and thyrotropes) at the expense of the more dorsal somatotropes and lactotropes (20,21). To determine if these same changes occur in heterozygous mutants, we analyzed the pituitary glands of newborn mice of all three genotypes, comparing heterozygous mutants to both wild-type and homozygous mutants. Since FSH does not terminally differentiate and is a cyclical hormone, our results for FSH were not interpretable. Therefore, we focused our analysis on 5 of the 6 anterior pituitary hormone cell types for neonates: ACTH, GH, PRL, TSH, and LH.

To quantify the relative numbers of each of these hormone-producing cell types in the anterior pituitaries of *Pax6^+/+^*, *Pax6^+/−^* and *Pax6^−/−^* neonates were serially sectioned in their entirety and separately immunolabeled for ACTH, GH, PRL, TSH or LH (Figure 3). The total numbers of specifically immunolabeled cells for each hormone were systematically counted in every other section so that for any given experiment about half the total number of cells in each anterior pituitary were assessed with respect to hormone cell type. The average number and standard deviation (SD) of immunolabeled cells per genotype were determined using three or four animals per group (Figure 3 D, H, L, P and T). Statistical comparison between genotypes was performed by analysis of variance (ANOVA). There were no significant changes in ACTH-cell numbers in either the heterozygote mutant or the homozygote mutant compared to wild-type littermates (Figure 3A-D). For neonates at P0, wild-type pituitary glands had an average of 3,622±444 ACTH expressing cells while *Pax6^+/−^* and *Pax6^−/−^* glands had averages of 4,217±380 and 3,964±962 expressing cells, respectively (Figure 3D). For PRL expressing cells, wild-type pituitary glands averaged 6485±992, *Pax6^+/−^* 7843±941, and *Pax6^−/−^* 8825±774, showing statistically significant difference between homozygous mutants and wild-type, but no statistically significant difference between heterozygous mutants and wild-type, although the trend suggests heterozygous mutants have increased lactotrope cell numbers (Figure 3 I-L). Significant changes in the numbers of GH- and TSH-expressing cells in the neonatal pituitaries did exist between the different genotypes (Figure 3H,P). The numbers of GH-expressing cells was decreased in both homozygous (5,574±1442 cells) and heterozygous (6,959±1103 cells) mutant genotypes compared to wild-type littermates (9,225±1049 cells). Conversely, the numbers of TSH-expressing cells was increased in both homozygous (5,854±810 cells) and heterozygous (4,828±817 cells) mutant genotypes compared to wild-type littermates (2,803±917 cells). The numbers of LH-expressing cells were generally numerically increased in both *Pax6^−/−^* (2,965±731 cells) and *Pax6^+/−^* (2,400±1448 cells) animals compared to wild-type (1883±372 cells), but the differences were not statistically significant (p = 0.44), largely due to the variance observed in the heterozygous samples (Figure 3T).

**Figure 3:**
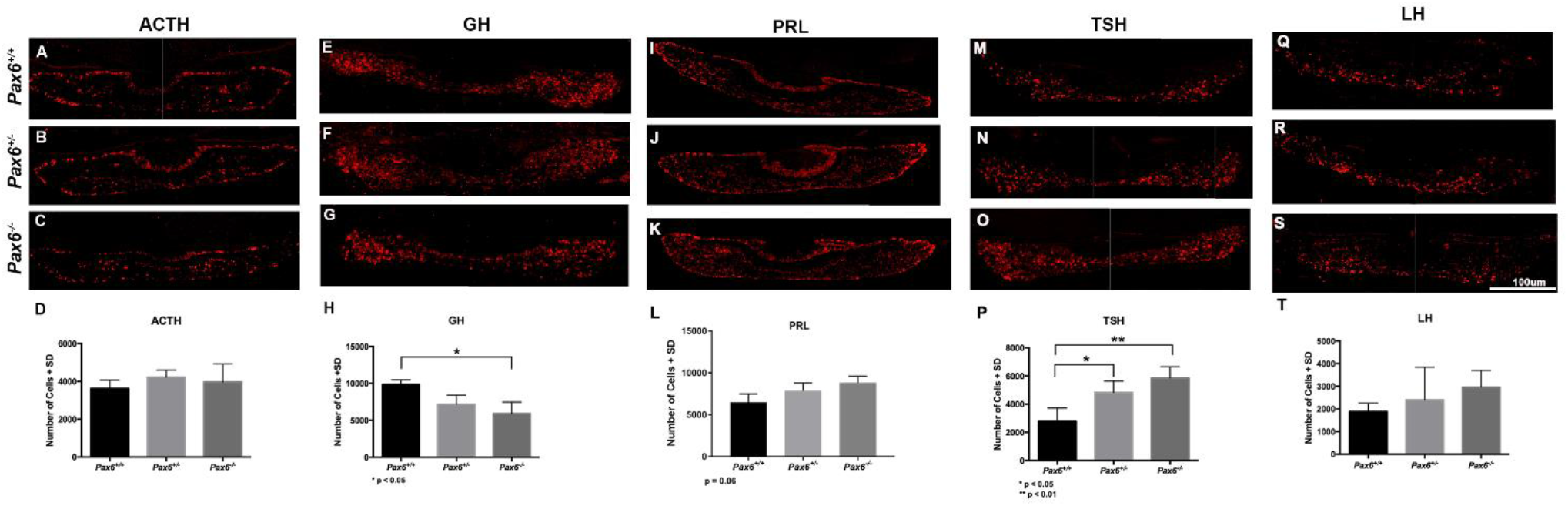
Reduction in *Pax6* results in an increase in TSH-producing cells in neonate mouse pituitary glands. Representative pituitary tissue sections from wild-type (A,E,I,M,G), *Pax6^+/−^* (B,F,J,N,R) and *Pax6^−/−^* (C,G,K,O,S) P0 neonates showing expression of ACTH (A-C), GH (E-G), PRL (I-K), TSH (M-O), and LH (Q-S). Quantification and comparison of the relative numbers of ACTH (D), GH (H), PRL (L), TSH (P) and LH (T) expressing cells in the pituitary of each genotype reveals that there is both an increase in TSH producing cells and a decrease in GH producing cells in both *Pax6^−/−^* and *Pax6^+/−^* mutant mice relative to wild-type. *p<0.05, **p<0.01. Data represented as mean ± std dev, n=3 for ACTH and LH, n=4 for PRL, TSH and GH for each genotype.

The comparative results for ACTH-, GH- and TSH-expressing cells between wild-type and homozygous mutant neonates are consistent with those reported for the embryonic pituitary (20,21). Interestingly, there are also differences in both GH- and TSH-expressing cell numbers in heterozygotes compared to wild-type, and the trends observed across genotypes suggests that the effect is *Pax6* dosage sensitive. Our results for LH-expressing cells appears to be inconsistent with the report by Kioussi et al. that there was a significant increase in the numbers of LH-expressing cells in homozygous mutant embryos relative to wild-type, but it is consistent with the LH data shown in the study by Bentley et al (21). The apparent discrepancies for this cell type may be due differences in the markers used (LH here and by Bentley et al; SF-1 by Kioussi et al), problems of sample size, or potential modifier effects between the mice used in these studies. Our results for PRL were also inconsistent with the reported decrease in PRL expressing cells by Kioussi et al. Similar explanations for those listed for LH might also explain the differences in results.

### Endocrine cell numbers in adult pituitary glands

To better understand the effect of *Pax6* haploinsufficiency on pituitary function in adult mammals, flow cytometry was used to quantify the relative numbers of ACTH, GH, PRL, TSH, FSH or LH hormone-producing cell types in the anterior pituitaries of adult *Pax6^+/+^* and *Pax6^+/−^* mice (6-8 months of age). There was no significant difference in the numbers of ACTH-, PRL-, GH-, FSH-, or LH-expressing cells between wild-type and heterozygous mutant mice (Figure 4). On average, 13% of cells positively stained for ACTH in wild-type pituitary glands and 11% in *Pax6^+/−^* pituitary glands; 28% for PRL in wild-type pituitary glands compared to 33% for *Pax6^+/−^* mice; 34% of cells positively stained for GH in wild-type glands and 31% in those of *Pax6^+/−^* mice; for FSH, 30% were positively stained in wild-type and 30% in *Pax6^+/−^* mice; and 9.6% positively stained for LH in wild-type mice compared to 7% in mutants. In contrast, the numbers of TSH-producing cells were significantly increased in *Pax6^+/−^* pituitaries compared to wild-type (Figure 4 P-T). On average, 19% of cells positively labeled for TSH in *Pax6^+/−^* mice compared to 11% in wild-type.

**Figure 4:**
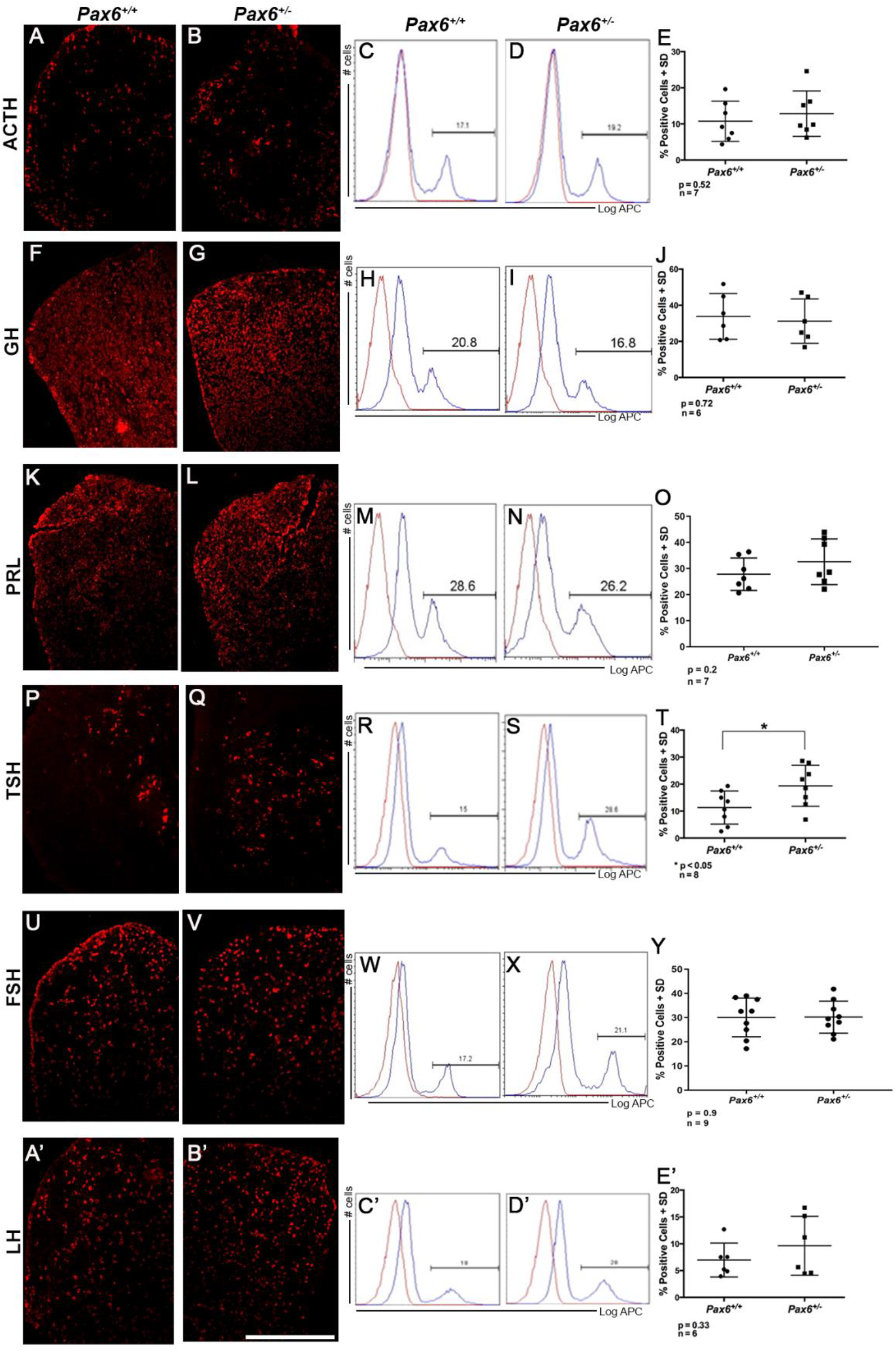
Increase in TSH-producing cells in *Pax6^+/−^* mice persists in adult pituitary glands. Representative pituitary tissue sections from wild-type (A,F,K,P,U,A’) and *Pax6^+/−^* (B,G,L,Q,V,B’) adult mice showing expression of ACTH (A,B), GH (F,G), PRL (K,L), TSH (P,Q), FSH (U,V) and LH (A’,B’). Representative results from flow cytometry performed on pituitaries from wild-type (C,H,M,R) and *Pax6^+/−^* (D,I,N,S) adults sorted for ACTH (C,D), GH (H,I), PRL (M,N), TSH (R,S), FSH (W,X), and LH (C’,D’). Quantification and comparison of the relative numbers of ACTH (E), GH (j), PRL (O), TSH (T), FSH (Y) and LH (E’) expressing cells reveals that there is an increase in the numbers of TSH-expressing cells in the pituitaries of *Pax6^+/−^* animals compared to wild-type. Data represented as mean ± std dev. Scale bar = 100um. Mice were 6-8 months old and include both sexes.

To assess the amount of protein present within the whole adult pituitary gland, we performed western blots on homogenized pituitary glands from *Pax6^+/−^* pituitary glands compared to wild-type littermates. For GH and LH, the amount of protein was equal between both genotypes (data not shown). The amount of TSH protein present was increased in *Pax6^+/−^* compared to wild-type, most likely a consequence of the increase in thyrotrope cell numbers in mutant pituitary glands. Interestingly, although there was no significant change in ACTH and PRL-expressing cell numbers, both of these hormones showed an increase in amount of protein in *Pax6^+/−^* pituitary glands (data not shown). This suggests that although *Pax6* does not change cell numbers of these hormone cell types, it may impact the processing and secretion of the hormone.

### Assessing the effects of increased number of TSH-producing cells in *Pax6^+/−^* mice

A possible effect of the increase in TSH-producing cells in the pituitaries of *Pax6* heterozygous mutant mice is a change in metabolism. This intrigued us because many patients with aniridia self-report difficulty managing their weight (16). Therefore, we first decided to measure the weights of *Pax6^+/−^* mice compared to their wild-type littermates. The mice were between 6 to 9 months of age and housed under directly comparable conditions. We found that as a group *Pax6^+/−^* mice weighed less than wild-type for both males and females. Weights for wild-type males averaged 46.0 ± 6.8g compared to average weight of 43.6 ± 6.8g for *Pax6^+/−^* mice (Figure 5A, n= 76 *Pax6^+/+^* and 76 for *Pax6^+/−^,* p = 0.03). Similarly, wild-type females weighed an average of 36.5 ± 7.0g and *Pax6^+/−^* females weighed an average 33.7 ± 7.0 (Figure 5B, n= 79 *Pax6^+/+^* and 79 for *Pax6^+/−^*, p = 0.01). Interestingly, there was a larger variation in the weights observed for both male and female *Pax6^+/−^* mice compared to their wild-type littermates (Figure 5A,B). These data suggest that metabolic control mechanisms and/or tolerances may be altered in these *Pax6^+/−^* mice compared to wild-type animals.

**Figure 5:**
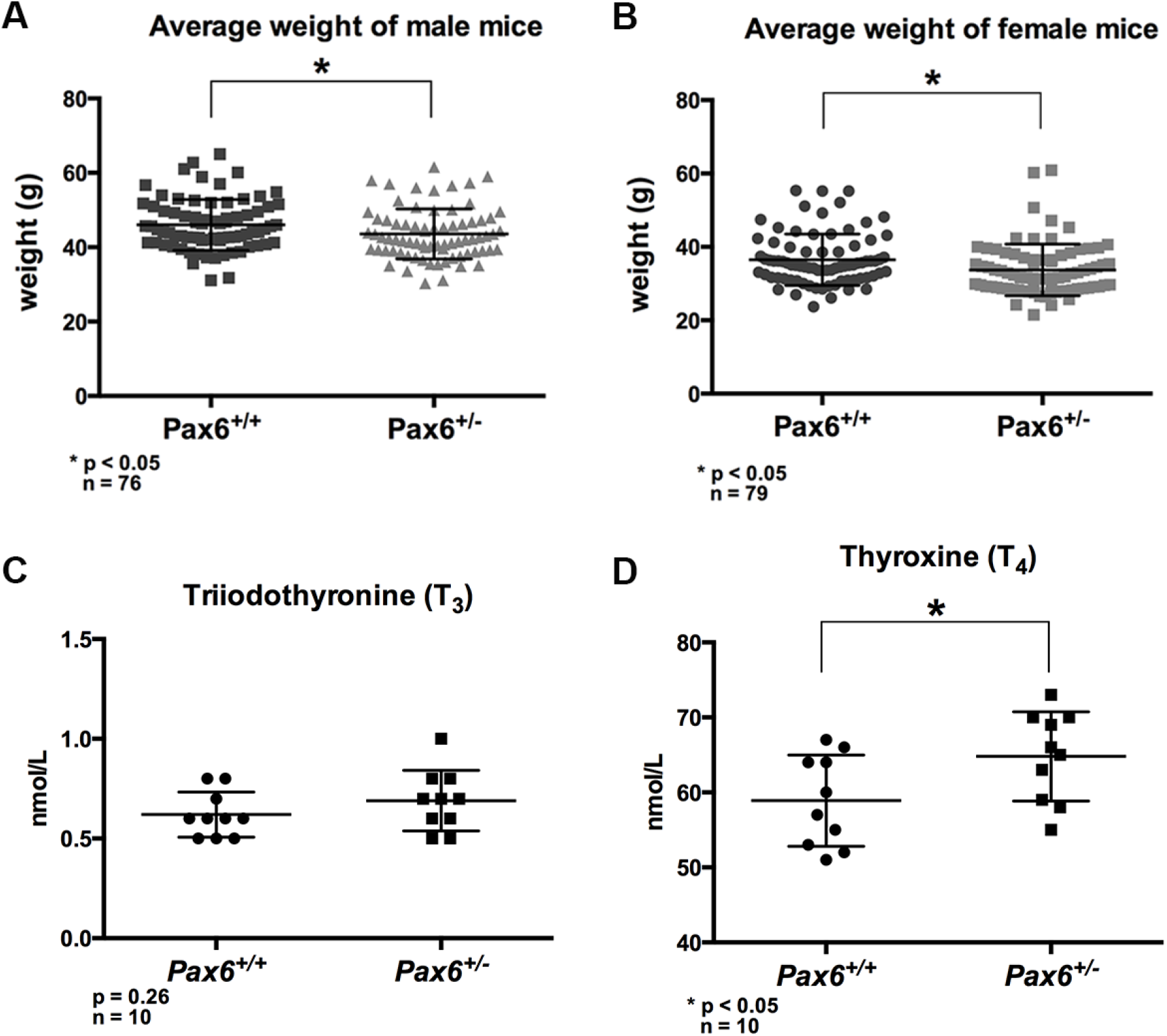
Adult *Pax6^+/−^* mice weigh less than wild-type littermates and have increased T_4_ levels in serum. (A) In both male and (B) female mice, *Pax6^+/−^* mutant mice collectively weigh less than their wild-type littermates. Mice were 6 to 9 months of age. For each genotype 76 male mice and 79 female mice were weighed. Range: male wild-type, 31.8-65.0 g; male *Pax6^+/−^*, 31.1-61.6 g; female wild-type, 23.7-55.0 g; female *Pax6^+/−^*, 21.0-60.0 g (C) Concentration of total T_3_ (TT_3_) measured reveals no significant difference whereas (D) concentration of total T_4_ (TT_4_) is significantly higher in *Pax6^+/−^* mice than in wild-type littermates. Mice were 4 months of age. Serum was collected from 6 males and 4 females per genotype. Range TT_3_ (males + females): wild-type, 0.50-0.80 nmol/L; *Pax6^+/−^*, 0.50-1.00 nmol/L. Range TT_4_ (males+females): wild-type, 51.0-67.0 nmol/L; *Pax6^+/−^*, 55.0-73.0 nmol/L. Data represented as mean ± 1 SD, *p<0.05

Given our finding that TSH cell numbers were increased in *Pax6^+/−^* mice and that thyroid hormones are known to be involved in the regulation of basal metabolism and thermogenesis (37–40), we wanted to assess the levels of TSH and thyroid hormones in these mice. TSH acts on the thyroid gland to stimulate release of triiodothyronine (T_3_) and thyroxine (T_4_). Because access to a validated TSH assay for mice proved problematic, we report only on our findings for circulating levels of T_3_ and T_4_, which also can provide a readout of TSH levels. The average concentration of T_3_ in *Pax6^+/−^* mice was 0.7 ± 0.2 nmol/L, slightly higher than the average concentration for wild-type of 0.6 ± 0.1 nmol/L, although this difference was not statistically significant (Figure 5C, p = 0.26). For T_4_, the average concentration for *Pax6^+/−^* mice was 65 ± 6 nmol/L, significantly higher than the average concentration for wild-type of 59 ± 6 nmol/L (Figure 5D, p = 0.04). Although there is no notable difference in levels of T_3_, the elevated concentration of T_4_ in *Pax6^+/−^* mice may lead to an increased metabolism, which may explain why our heterozygous mutant mice weigh significantly less than their wild-type littermates.

## Discussion

Recent studies have suggested that adult patients with aniridia have a greater propensity for unexplained systemic symptoms, such as infertility, severe eczema and issues with weight, which may be related to *PAX6* haploinsufficiency (16). The goal of this study was to test if any changes were present in the adult anterior pituitary gland as a result of a heterozygous mutation in the *Pax6* gene in mice and, if so, whether or not these changes could explain some of the symptoms experienced by patients with aniridia.

This study represents the first report of *Pax6* expression and function in the adult pituitary gland. We found that *Pax6* expression is maintained in the anterior gland in adult mice, and expression is no longer restricted to the dorsal region, but is expressed throughout the anterior lobe much like it is at E14.5. This suggests that the maintenance of *Pax6* expression may be required for proper functioning of the anterior pituitary as it is in the pancreas. In the developing pancreas, cells of the endocrine lineage express *Pax6,* and a complete loss of *Pax6* results in both lack of development of α-cells and a marked decrease of all other cells types in the endocrine pancreas (41). In the adult, *Pax6* is expressed throughout the islets of Langerhans and continued expression is required for normal pancreatic function (4). In the case of the pituitary, *Pax6* is not required for specification of the hormone producing cells, as evidenced by the fact that in the complete absence of *Pax6* all cell types still develop, but *Pax6* is required for generation of normal numbers of somatotropes, lactotropes and thyrotropes (current study (20,21)), and could play a role in the ability of the adult pituitary to adjust to physiological demand. This could occur at the level of endocrine cell function or in mediating changes in cell numbers. Changes in physiological demand have been shown to result in changes the populations of hormone-producing cells in the adult pituitary, and this can occur through proliferation of terminally differentiated cells; transdifferentiation of differentiated cells, such as conversion of somatotrophs to lactotrophs; and/or differentiation of progenitors/stem cells (42). In the nervous system, *Pax6* is known to regulate potency (43), cell-cycle and differentiation kinetics (44–50), the balance between neural stem cell self-renewal and neurogenesis (51), and is required to properly respond to signals during lens induction (52). In the context of endocrine cell function, *Pax6* is implicated in the normal function of intestinal L cells by direct activation of the *glucagon~like peptide 1* gene (5–7), and of islet cells by regulation of a number of genes required for endocrine cell function, including the production of insulin, glucagon and somatostatin (23,53–57). Thus, it is plausible that a reduction in *Pax6* protein levels could affect pituitary function in adult mammals.

Significantly, we found an increase in the numbers of TSH-producing cells in both neonatal and adult mice heterozygous null for *Pax6*, and this correlated with a significant increase in TSH protein from *Pax6* heterozygous null pituitary glands and in circulating levels of T_4_ in adult animals. An increase of T_4_ levels in the serum above normal was consistent with a putative increase in the amount of TSH being produced by the pituitaries in *Pax6^+/−^* animals. When stimulated by TSH, the thyroid gland releases T_4_ and T_3_ into the blood (58–65). An increase in circulating levels of T_4_ would be expected to also lead to a concomitant increase in the levels of T_3_ in the brain, liver, kidney, intestines and other target tissues (63,66–75). In rodents and other mammals, T_3_ is estimated to be 3 to 5 times more potent than T_4_ (76–79). Thus, a modest increase in circulating T_4_ levels could result in a significant change in the metabolic potency of T_4_ in target tissues due to its conversion to T_3_.

While the physiological effects of abnormally high levels of circulating T_4_ in *Pax6^+/−^* mice were not fully assessed, one possibility was that these animals were exhibiting symptoms associated with mild hyperthyroidism. Such changes could include increased appetite, increased metabolism, tachycardia, heat intolerance, fatigue, and anxiety and irritability. Consistent with this hypothesis, we found that both male and female *Pax6^+/−^* adult mice weighed less than their wild-type littermates housed under identical conditions. Similar results were obtained in two other studies examining the role of *Pax6* in metabolic homeostasis (7,80), and the authors noted that although adult *Pax6^+/−^* mice showed increased food intake compared with wild-type control mice, they were resistant to diet-induced fat accumulation (7).

It may be difficult, however, to disentangle changes specifically associated with elevated levels of T_4_ from *Pax6* mediated changes in other metabolic systems. In addition to its role in the development and function of the eye and brain, *Pax6* is known to be required for normal development and function of the endocrine pancreas (4,41), and it has been implicated in the normal development and/ or function of the enteroendocrine cells that line the intestine (5–7). Given that metabolic pathways are tightly regulated, and that changes in *Pax6* can affect the development and/or function of the pituitary (current study, (20,21)), pineal gland (81,82), pancreas (4,41,56,83–86), enteroendocrine L-cells (5–7), and neuroendocrine neurons in the hypothalamus (23), it is likely that the physiological effects of increased thyroid hormone activity will act synergistically and/or additively to *Pax6*-dependent changes in these other metabolic pathways. Two tissues that bear closer examination are the liver and the kidney, where chronically elevated levels of thyroid hormones can result in clinically important alterations the function of these organs (87,88). In addition to possible effects in the adult, elevated levels of T_4_ could impact cortical development directly (89,90).

Although we found that as a group *Pax6^+/−^* adult mice weighed less than *Pax6* normal mice, individuals with aniridia report a higher incidence of obesity as adults (16). While this apparent discrepancy could be due to inherent metabolic differences between rodents and humans, it could also reflect changes in the metabolic control mechanisms and/or tolerances associated with a reduction in functional *Pax6* protein. Consistent with this idea, we noted a larger variation in the weights observed for both male and female *Pax6^+/−^* mice compared to their wild-type littermates, and some of the female *Pax6^+/−^* mice were much heavier than wild-type littermates (Figure 5B). Thus, while *Pax6^+/−^* mice maintained on normal rodent chow exhibited lower body weights as a group, these animals may exhibit a higher BMI compared to wild-type animals when provided access to a calorically rich diet. In a recent report, *Pax6^+/−^* mice fed a high fat diet gained more weight than wild-type control mice (Figure 1A in (80)); these mice also exhibited insulin resistance, reduced prohormone convertase 1/3 production, and increased proinsulin secretion (80). A possible link between *Pax6* function and obesity in the general population has been reported (91). It is important to note that not all *Pax6^+/−^* mice or persons with aniridia exhibit a high BMI, and that in general both juvenile mice and children with aniridia do not appear to be overweight (Johnson and Lauderdale, unpublished).

In addition to an increase in TSH-labeled cells, *Pax6^+/−^* neonates exhibited a reduction in the number of GH-labeled cells in the pituitary. We initially thought that the increase in TSH-producing cells was at the expense of GH-producing cells, consistent with the model proposed by Kioussi *et al*. (20). However, this seems unlikely since the number of GH-producing cells increased to normal in adult *Pax6^+/−^* mice, suggesting that GH cell fate determination was delayed or subject to a compensatory mechanism. Consistent with this idea, Bentley, *et al*. (21) found that at E17.5 GH serum levels were five-fold higher in wild-type embryos compared to *Pax6* null embryos, but that this difference was reduced to three-fold by P0. During normal pituitary organogenesis, there is an increase after birth in the numbers of the different hormone producing cell types and this expansion is driven by the hypothalamic releasing hormones and by physiological demands (42). In the case of somatotropes, this increase involves reentry into the cell cycle (92), and interestingly, requires thyroid hormone. Although young adult mice deficient in thyroid hormone production exhibit a 4-fold decrease in the numbers of GH-expressing cells in their pituitaries, treatment of these animals with T_4_ from birth increases GH cell numbers to those expected in normal age-matched animals (93). While the mechanism underlying the recovery of normal numbers of GH-expressing cells in the adult pituitary of *Pax6^+/−^* mice is not known, it may be driven in part by elevated levels of T_4_. Regardless of the mechanism, the recovery of GH cells would explain why *Pax6^+/−^* mice do not show any of the physical attributes associated with a decrease in GH.

Three recent studies have examined pituitary function in aniridia patients with heterozygous *PAX6* mutations (22–24). The first study performed an endocrinological evaluation on related individuals with aniridia, which consisted of a family with two affected individuals and a second family with 36 members, distributed in five generations, of whom 14 members were affected (22). These individuals were tested for FSH, LH, ACTH, prolactin, GH, somatomedin C, TSH, cortisol, estradiol concentrations, synacthene, and luteinizing hormone-releasing hormone (LHRH). Of these individuals, endocrine changes were detected in one person with aniridia from the first family (low ACTH concentration) and in two individuals with aniridia in the second family (reduced response to synacthen/ACTH test). The second study examined a pedigree with 19 individuals with aniridia spread over three generations and reported that the levels of ACTH (the only pituitary hormone assessed) and α-MSH were decreased in the blood of the aniridia patients as a group (23). The third study reported a single case, which was that of a 40 year old women who had obesity, exhibited an impaired glucose tolerance with a delayed insulin response, and also exhibited slightly impaired pituitary function, which consisted of subclinical hypogonadotropic hypogonadism and borderline GH deficiency, but apparently normal ACTH levels (24).

One conclusion that can be drawn from these studies is that individuals with *PAX6*-mediated aniridia do not necessarily exhibit overt clinical symptoms associated with abnormal pituitary function. The reduction in ACTH levels can be attributed to a PAX6-dependent reduction in the levels of prohormone convertase (PC1/3) expressed in the hypothalamus (23). PC1/3 is the enzyme responsive for converting pro-opiomelanocortin (POMC) into ACTH, which is further converted into α-MSH (94). In the case of the single individual with aniridia exhibiting somewhat reduced levels in FSH, LH, and GH, but apparently normal levels of ACTH, the changes could be explained by a mechanism unrelated to *PAX6* or may reflect phenotypic variation.

When compared to our findings in mice, it is possible that *Pax6* haploinsufficiency affects the human pituitary differently than in rodents; however, we think it likely that *Pax6* haploinsufficiency affects mammalian pituitaries in similar ways. Our studies show that, in mice, a loss of function mutation in *Pax6* results in increased numbers of TSH-producing cells in the developing and adult pituitary gland, leading to an increase in circulating concentration of total T_4_ as measured in a fairly homogeneous group of mice. It is possible that there are comparable changes in persons with aniridia, but that there is more phenotypic variability. As described above, background-dependent variation particularly in somatotropes and lactotropes, but also corticotropes and thyrotropes, have been observed in mice (21). In the case of ACTH, although we did not find a change in corticotrope cell numbers between *Pax6^+/−^* and wild-type animals, *Pax6^+/−^* mice are reported to have a reduction of ACTH and α-MSH in hypothalamic tissue similar to humans with aniridia (23).

In conclusion, several lines of evidence indicate that the *Pax6* gene plays a role in the development and function of the endocrine system, and that the normal function of this system can be perturbed by *Pax6*-haploinsufficiency. Continued investigation into this important question is needed if we are to understand the systemic aspects of aniridia.

## Abbreviations

ACTH: adrenocorticotropin hormone;
CSF: cerebro-spinal fluid;
FSH: follicle stimulating hormone;
GH: growth hormone;
H&E: hematoxylin and eosin stain;
ISH: in situ hybridization;
LH: leutenizing hormone;
MRI: magnetic resonance imaging;
P: pituitary;
PRL: prolactin;
TSH: thyroid stimulating hormone;
V: third ventricle;
VBM: voxel-based morphometry

## Acknowledgments

The authors thank Drs. Sally Camper and Peter Gergics (Department of Human Genetics, University of Michigan Health System) and Dr. Shannon Davis (Department of Biological Sciences, University of South Carolina) for helpful discussions and technical advice on tissue preparation of neonate pituitary glands and immunolabeling cells in the pituitary, and Dr. Julie Gordon (Department of Genetics, University of Georgia) for helpful discussion and comments on the manuscript. Confocal microscopy was performed using the services of the Biomedical Microscopy Core under the direction of Dr. Muthugapatti K. Kandasamy. Flow cytometry was performed using the facilities of the CTEGD Cytometry Shared Resource Lab under the direction of Ms. Julie Nelson. The authors thank A. F. Parlow at the National Hormone and Peptide Program (NHPP), Harbor-UCLA Medical Center (Torrance, CA 90509, USA), for providing antibodies used in this study, and the Michigan State University Diagnostic Center for Population and Animal Health (Lansing, MI 48909-7576, USA) for assessing serum levels of T3 and T4.

